# Ureteric Bud Cells Programmed from Embryonic Stem Cells Obtain Competence for Secondary Induction in the Kidney

**DOI:** 10.1101/722157

**Authors:** Zenglai Tan, Aleksandra Rak-Raszewska, Ilya Skovorodkin, Seppo J. Vainio

## Abstract

Generation of kidney organoids from pluripotent stem cells (PSCs) is regarded as a potentially powerful way to study kidney development, disease, and regeneration. Direct differentiation of PSCs towards renal lineages is well studied, however, most of the studies relates to generation of nephron progenitor population from PSCs. Until now, differentiation of PSCs into ureteric bud (UB) progenitor cells demonstrates limited success. Here, we describe a simple, efficient and reproductive protocol to direct differentiation of mouse embryonic stem cells (mESCs) into UB progenitor cells. The mESC–derived UB cells were able to induce nephrogenesis when placed in the interaction with the primary metanephric mesenchyme (pMM). In generated kidney organoids, the embryonic pMM developed nephron structures and the mESC-derived UB cells formed network of collecting ducts, connected with the nephron tubules. Altogether, our studies established an uncomplicated and reproducible platform for kidney disease modelling, drug testing and regenerative medicine applications.

## INTRODUCTION

Pluripotent stem cells (PSCs) possess great potential of differentiating into multiple cell types that are wildly used for developmental studies and regenerative medicine (Daisy and George, 2012). The kidney organoids derived from PSCs have been shown to be able to mimic the *in vivo* kidney structures development and function *in vitro* (Takasato et al., 2015, Morizane et al., 2015). The renal organoids in the four-dimensional (4D) (3D plus time) culture system self-organize highly complex tissue specific morphology, that are sufficient to model tissue development, disease and injury (Boreström et al., 2018, Kim et al., 2017, Taguchi and Nishinakamura, 2017, Lorna et al., 2018). The *in vitro* kidney organoid system provides new regenerative strategies as the combination of genome editing and stem cell technologies allow to generate personalized kidney organoids, which provide powerful tool for kidney disease treatment and drug toxicity screening (Czerniecki et al., 2018).

Recently, multiple reports presented protocols enabling generation of renal lineages from mouse and human PSCs. We and several other groups reported induction of nephron progenitors, which have the potential to develop into epithelial nephron like structures (Freedman et al., 2015, Garreta et al., 2019, Homan et al., 2019, Morizane et al., 2015, Taguchi et al., 2014, Takasato et al., 2014, Takasato et al., 2015, Tan et al., 2018, Low et al., 2019). Other groups have shown the derivation of ureteric bud (UB) progenitors, however, they didn’t show nephron progenitor induction (Xia et al., 2013) and also lacked the connection between collecting ducts and nephrons (Xia et al., 2013, Takasato et al., 2015). A newly published study have shown the generation of UB structures from PSCs, which possessed UB-like branching morphogenesis when aggregated with the embryonic metanephric mesenchyme (MM) to form a chimeric kidney organoids (Taguchi and Nishinakamura, 2017). However, these organoids lacked vascularization and the protocol is technically complex which limits its application. Analysis of these reports suggest, that we still need further study to develop simple, reproductive and stable protocols of UB progenitors generation.

Here, we report a simple protocol to direct differentiation of mouse embryonic stem cells (mESCs) into UB progenitor cells. The generated UB progenitor cells own the potential to develop into mature ureteric bud structures and have the capacity to induce nephrogenesis when co-cultured with dissociated pMM. In the reconstructed kidney organoids, the pMM developed into nephron structures and the UB progenitor cells formed the collecting ducts, which also connected with the nephron tubules. The chimeric kidney organoids also display presence of endothelial cells forming vascular network. Moreover, we have also shown that this chimera organoid setting can be used to study mechanisms of kidney development and drugs toxicity. In conclusion, our studies established an uncomplicated and reproducible method to generate UB progenitors from PSCs which can be used for tubulogenesis induction, and further for kidney disease modelling and regenerative medicine applications.

## RESULTS

### Direct differentiation of mESCs into UB progenitor cells

Both, the nephron and ureteric bud progenitor cells are derived from the intermediate mesoderm (IM). To establish a protocol and direct differentiation of mESCs into UB lineage, we first differentiated mESCs into IM (Figure 1A). We treated the mESCs with FGF2 and activin A to differentiate mESCs into epiblast in monolayer culture (Figure S1). The differentiated cells showed the expression of epiblast markers like *Fgf5* and *T (Brachyury)* (Figure 1B). We then used GSK-3β inhibitor, the CHIR99021 (CHIR), together with low concentration of BMP singling inhibitor, the Noggin, to activate mouse epiblast cells to primitive streak (PS), these cells expressed PS markers: *Cdx2*, T, *Tbx6* and *Mixl1* (Figure 1B). We followed for 2 days with activin A treatment, and the cells differentiated to IM stage and showed the expression of *Osr1, Lhx1* and *Pax2* (Figure 1B).

**Figure 1.**
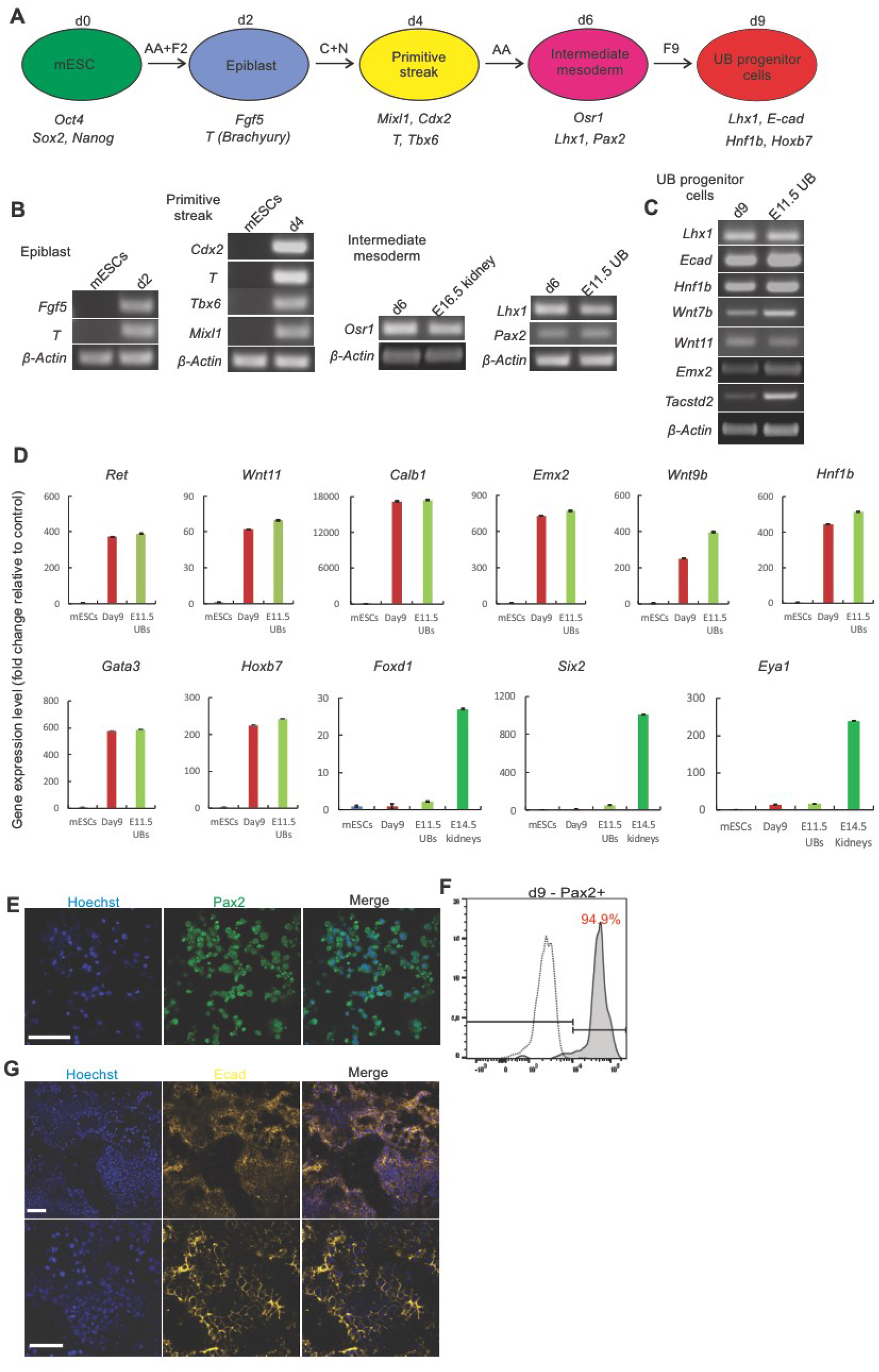
Differentiation of mouse ESCs to UB progenitor cells. (A) Schematic of the differentiation protocol of mESCs into UB progenitor cells. AA: Activin A; F2: FGF2; C: CHIR99021; N: Noggin; F9: FGF9. (B) RT-PCR displays mESCs differentiation into epiblast markers, primitive streak markers and intermediate mesoderm markers; *β-actin* was used as a housekeeping gene, the E16.5 kidney, E11.5 UB and mESC were used as a control. (C) RT–PCR at day 9 of differentiation showing the expression of markers of UB progenitors; the E11.5 UB was used as positive control. (D) qRT–PCR showing the gene expression [fold change] of ureteric bud markers; no expression of stroma (*Foxd1*) and nephron progenitor cell markers (*Six2* and *Eya1*) were observed at day 9 of differentiation. UBs, ureteric buds. (n = 3). (E) Immunocytochemistry and (F) flow cytometry of Pax2 in mESCs on day 9 of differentiation. Sample stained with secondary antibody alone was used as control for flow cytometry (dotted line). Scale bars, 50 μm (E). (G) Immunocytochemistry for E-cadherin (Ecad) on day 9 of differentiation; Scale bars, 50 μm.

Then we treated these cells with moderate concentration of FGF9 for additional 2-3 days, directing them to differentiate into the UB progenitor cells with expression of UB markers: *Lhx1, Ecad, Hnf1b, Wnt7b, Wnt9b, Ret, Calb1, Wnt11, Emx2, Gata3, Hoxb7* and *Tacstd2* (Figures 1C and 1D). In addition, no expression of stromal cell marker *Foxd1* and nephron progenitor cell markers *Six2* and *Eya1* (Figure 1D) at day 9 of differentiation was observed. The Pax2 immunostaining and FACS analysis showed that more than 90% of cells were Pax2+ (Figures 1E and 1F). Immunocytochemistry also revealed that use of moderate concentration of FGF9 induced the cells to express E-cadherin (Ecad) (Figure 1G) which may suggest that these differentiated cells represent putative UB progenitor cells.

### Generation of kidney organoids by mESC-derived UB progenitor cells and dissociated primary MM population

We and other groups previously reported that dissociation of mouse embryonic MM into single cells maintain the nephron progenitor stemness. The dissociated MM population develops into nephrons when induced by the inducer such as the embryonic UB or spinal cord (Saxén and Sariola, 1987, Junttila et al., 2015, Taguchi and Nishinakamura, 2017). To establish the potential and function of the mESC-derived UB progenitor cells, we aggregated these cells with mouse E11.5 dissociated pMM cells to generate a kidney organoid. We transferred the aggregated cell pellets into the traditional Trowell organ culture system. The cell aggregates were cultured for up to 11 days, during which they spontaneously formed kidney organoids with complex structures (Figures 2A and 2B). On day 3, we observed early Troma1+ UB formation, and formation of renal vesicle adjacent to the UB (Figure 2C).

**Figure 2.**
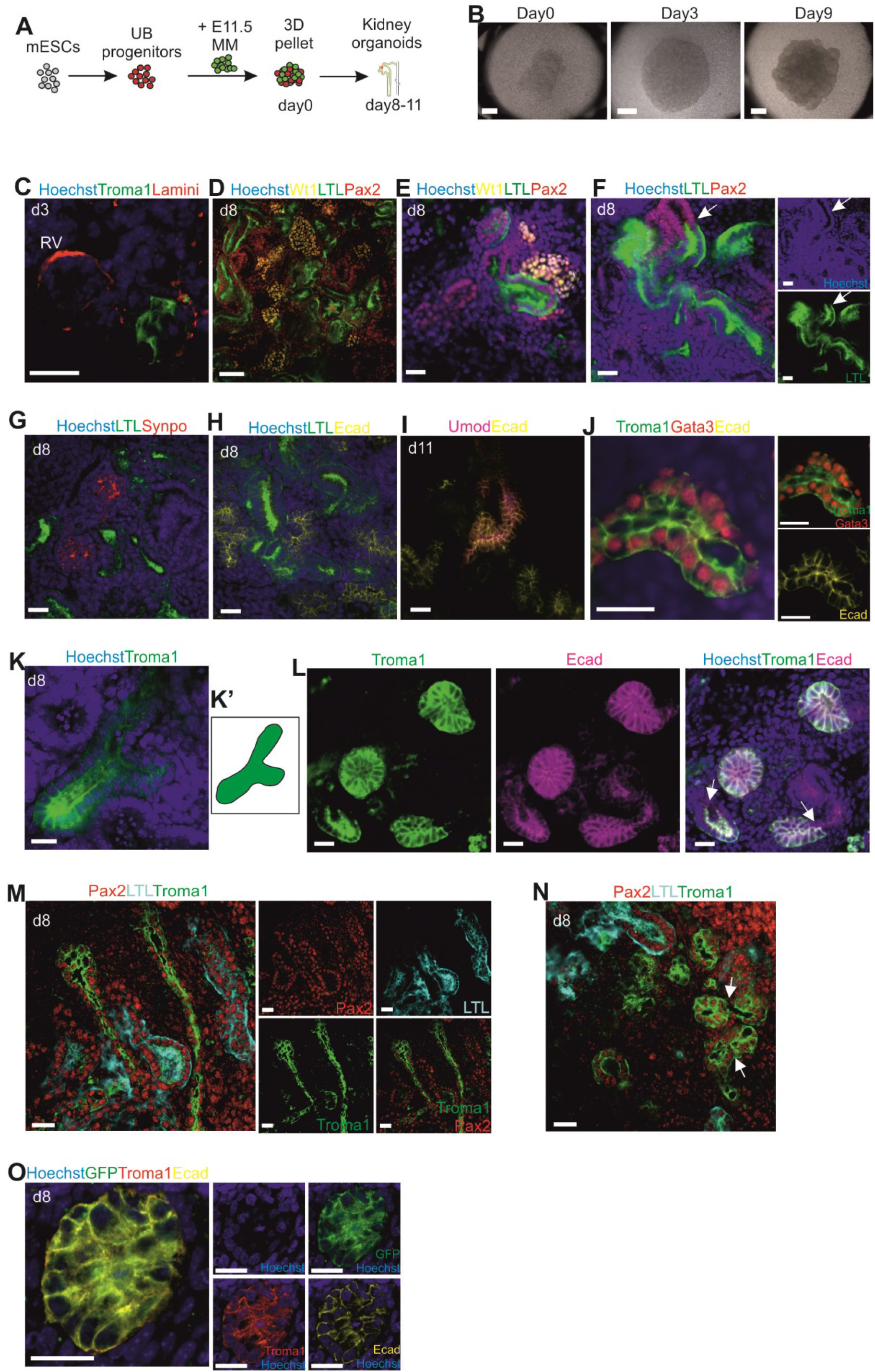
Generation of renal organoids by mESC-derived UB progenitors and pMM cells. (A) Schematic of generation of kidney organoids from mESC-derived UB progenitors with mouse E11.5 dissociated pMM. (B) Global bright field images of self-organizing kidney organoids in the Trowel organ culture. Scale bars, 500 μm. (C) Immunofluorescence of kidney organoids at day 3 show the formation of the renal vesicle next to the Troma1+ structure generated by mESC-derived UB progenitor cells. Scale bar, 20 μm. (D-M) Immunofluorescence of kidney organoids at either day 8 or 11. (D) Glomeruli are marked with Wt1, proximal tubules are marked with LTL and distal tubules are marked with Pax2+LTL-. (E) A single developing nephron; WT1+ glomeruli observed next to LTL+ proximal tubule. (F) Glomeruli (Synpo+) adjacent to proximal tubules (LTL+). (G) Proximal tubules (Pax2+LTL+) connected with distal tubules (Pax2+LTL-). (H) Confocal image shows Proximal tubules (LTL+) connected with distal tubules (Ecad+LTL-). (I) Loops of Henle marked by Umod and Ecad on day 11. (J) Confocal images of Troma1+Gata3+Ecad+ collecting duct structure. (K-K’) A “T” shaped UB structure in the kidney organoid. (L) mESC-derived UB progenitor cells generated collecting ducts (Troma1+Ecad+) connecting with nephron distal tubules (Troma1-Ecad+). (M) Kidney organoids developed collecting duct trunk structures. (N) The collecting ducts fused together. (O) Troma1+Ecad+ collecting ducts are derived from GFP+ mESCs. Scale bars, 50 μm (D), 20 μm (E-O).

The whole-mount immunostaining of day 8 chimeric organoids presented development of nephrons with the positive staining of glomeruli markers Wilms tumor 1 (Wt1+) and Synaptopodin (Synpo+); proximal tubule markers *Lotus tetragonolobus lectin* (LTL+), distal tubules markers Pax2+LTL- and Ecad+ (Figures 2D-2H, and S2A). Moreover, we found numerous Synpo+ and Wt1+ glomeruli adjacent to LTL+ proximal tubules, LTL+ proximal tubules connected with Pax2+LTL-/Ecad+ distal tubules (Figures 2D–2H and S2A), indicating a proper nephron structure: glomerulus/proximal tubule/distal tubule organization. On day11, the organoids also displayed Henle’s loop with the expression of markers uromodulin (Umod+) and Ecad+ (Figure 2I).

We also test whether the mESC-derived UB progenitor cells have the *in vivo* UB capacity, to form the collecting duct system following the kidney development. On day 8, by immunocytochemistry, we observed Troma1+Gata3+Ecad+Pax2+LTL-collecting duct structures in the kidney organoids (Figures 2J–2N and S2B and S2C). We also found some Troma1+ “T” shaped UB structures (Figures 2K-2K’). Importantly, we observed that the collecting ducts (Troma1+Ecad+) connected with the distal tubules (Ecad+Troma1-) of the nephron structure (Figures 2L and S2C) generating interconnection between collecting ducts and nephrons, so essential for urine drainage. The collecting ducts displayed the morphology of branches and long collecting duct trunks (Figure 2M). In addition, some of the collecting ducts fused together in one duct, similar to the *in vivo* collecting duct tree (Figure 2N and Video 1). Altogether, the *in vitro* reconstructed organoids developed towards kidney structures that are similar to *in vivo* kidney.

To identify, that the kidney organoid collecting duct structures were derived from mESCs, we have generated the mESC line with stable expression of the GFP. Immunofluorescence analysis demonstrating that the Troma1+Ecad+ collecting duct were derived from GFP+ mESCs (Figures 2O and S2D). Next, to rule out the possibility that the differentiated cells (mESC-derived UB progenitors) could give rise to nephrons when induced by the embryonic UB, we aggregate the day 9 differentiated cells with E11.5 dissociated UB cells and transferred them to organ culture. Since the UB survival and development depends on the presence of metanephric mesenchyme, in the organ cultures the purified UB failed to branch in the presence of UB-differentiated cells (Figure S2E). These data suggest that we did differentiate mESC towards UB progenitors, and they could further develop into collecting ducts when co-cultured with pMM.

We have also attempted to induce vascularization of developing glomeruli in these renal organoids. Previous studies showed that the ROCK kinases are VEGF downstream effectors, which negatively regulate the process of angiogenesis (Jens et al., 2009). Therefore, we used the ROCK inhibitor to enhance angiogenesis in the renal organoids. However, this treatment did not increased endothelial network area and the CD31+ cells could not be found in the developing glomerular tuft (Figure S3).

### Characterization of kidney organoids

Next, we wanted to verify, that the nephron structures in the organoids were generated via the MM population induced by the mESC-derived UB progenitors, and not by contamination of the MM with the primary UB cells. We cultured the E11.5 MM tissue in isolation and it underwent apoptosis already at the second day of culture and died at day 3 (Figure S4). This also confirmed that without suitable inducer the MM cells will not undergo nephrogenesis and cannot survive for long time in the organ culture condition in vitro.

To further confirm that the nephron structures and collecting ducts were derived from pMM and mESC-derived UB progenitors respectively, we used the mTmG (td-tomato) pMM population to aggregate with the differentiated mESCs (Figure 3A). The aggregates formed well-developed nephron structures (WT1+ glomeruli; LTL+ proximal tubules; Ecad+Troma1- distal tubules) which originated from the MM cells (mTmG+) (Figures 3B and 3C), and the Troma1 labeled collecting duct which was mTmG- and therefore originated from the mESC-derived UB progenitors (Figures 3C and 3D). These data presents that the kidney organoids are specifically derived from the interaction between the pMM and mESC-derived UB progenitors.

**Figure 3.**
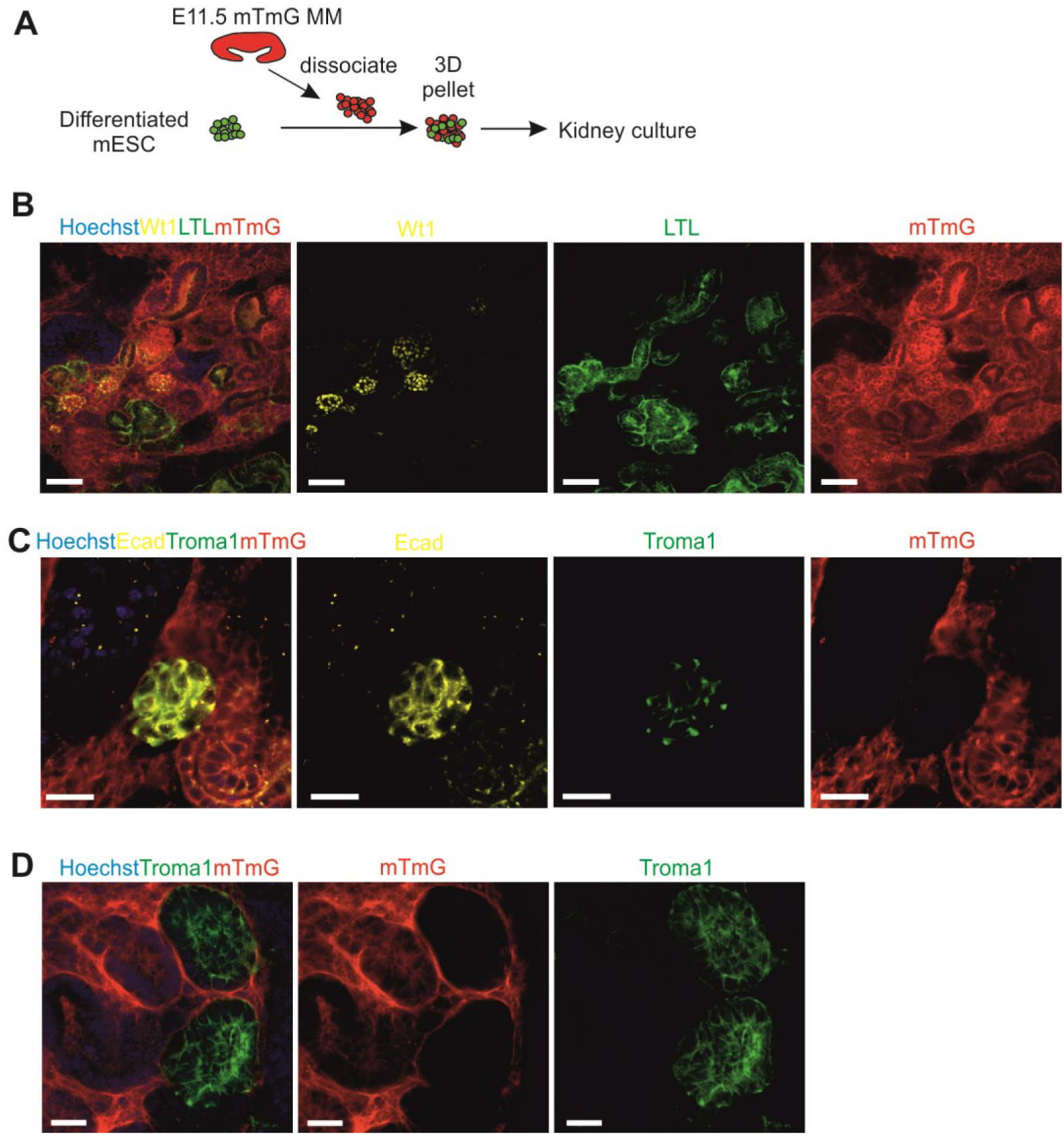
The UB structures in kidney organoids are specifically derived from differentiated mESCs. (A) schematic of generation of kidney organoids from mESC-derived UB progenitors with mouse E11.5 dissociated mTmG MM. (B) Confocal images of the 3D kidney organoids derived from E11.5 mTmG MM and mESC-derived UB progenitors. The nephron structures such as WT1+ glomeruli and LTL+ proximal tubules were derived from mTmG+ MM. Scale bars, 50 μm. (C) Confocal images of the kidney organoids showing the ureteric bud structures (Ecad+Troma1+) being generated by mESC-derived UB progenitor cells. Scale bars, 20 μm (D) The Troma1+ UB structures derived from mTmG-cells (mESC-derived UB progenitors). Scale bar: 20 μm.

### mESC-derived UB progenitor cells specifically integrated into the 3D ureteric bud structures

In order to identify the mESC-derived UB progenitors from wild type UB cells in chimeric organoids, and to ensure they will only generate UB cells, we have generated the organoid with the mESC-GFP line. We dissociated the whole E11.5 embryonic kidney rudiment (MM and UB) and mixed it with undifferentiated mESC or mESC-derived UB progenitors generating chimeric kidney organoids (Unbekandt and Davies, 2010, Junttila et al., 2015, Xia et al., 2013) (Figure 4A).

**Figure 4.**
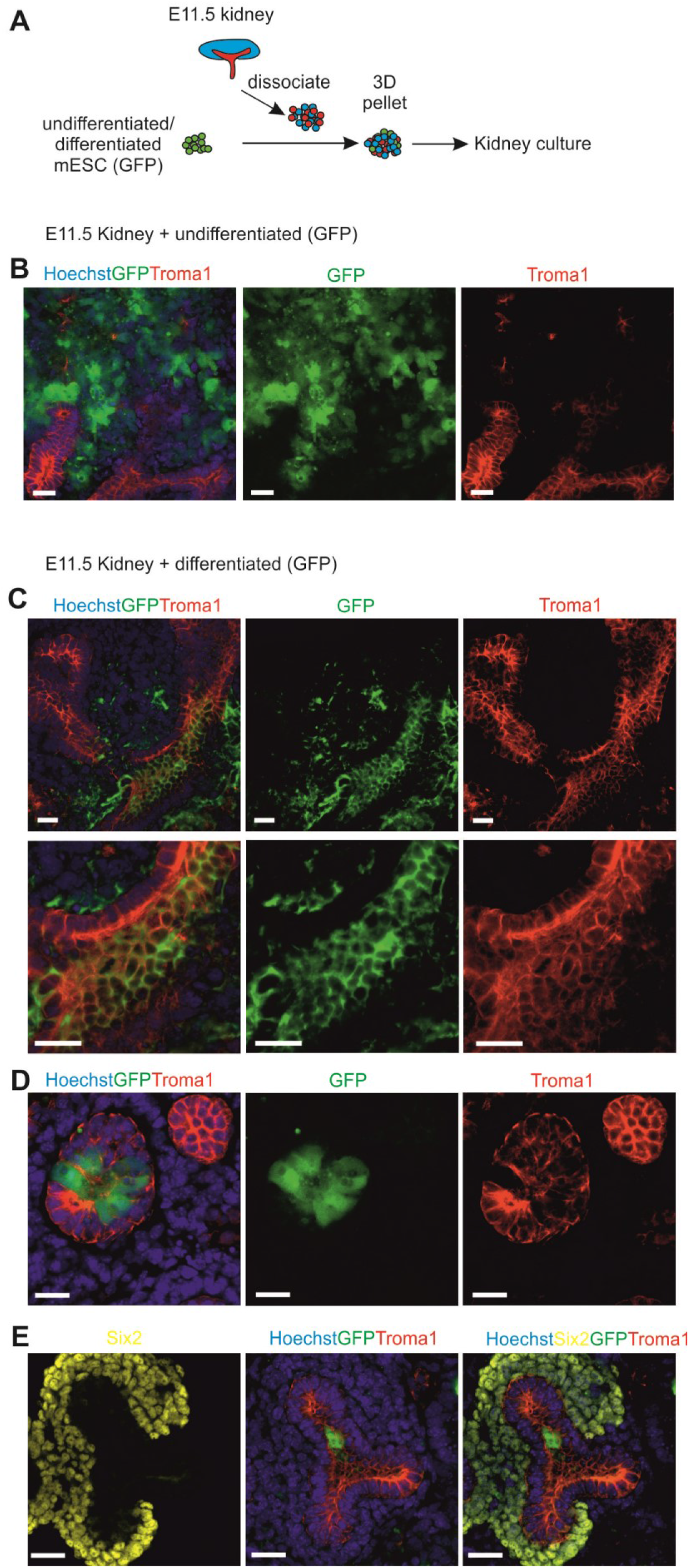
Mouse ESC-derived UB progenitor cells demonstrate a UB committed fate and integrated into embryonic UB on 3D organ culture *in vitro*. (A) Schematic of generation of kidney organoids from mESC-derived UB progenitors with mouse E11.5 dissociated kidney rudiments. (B) Immunofluorescence analysis demonstrating random localization of undifferentiated mESCs in organ co-cultures. Scale bars, 20 μm. (C) Immunofluorescence analysis demonstrating the localization and integration of the mESC-derived UB progenitors to the UB structures in chimeric organoids. Scale bars, 20 μm. (D) Immunofluorescence analysis demonstrating mESC-derived UB progenitors integrated into the UB structures and enhanced chimeric ureteric bud formation. Scale bars, 20 μm. (E) Confocal image presenting specific location of mESC-derived UB progenitor cells (GFP+) into Troma1+ UB structures of the chimeric organoid. Scale bar: 20 μm.

The undifferentiated mESCs aggregated with dissociated embryonic kidney rudiments presented random localization within the organoids and had negative effect on the nephrogenesis. The undifferentiated mESC had formed groups within the renal structures; however, the structures were not many and even only very few Troma1+ ureteric buds could be observed (Figure 4B).

When we aggregated the mESCs-derived UB progenitors with the dissociated embryonic kidney, we noticed much more robust tubulogenesis and numerous chimeric ureteric bud structures (Figures 4C and 4D). The UB progenitor cells did not interfered with renal development and efficiently and specifically integrated into the mouse ureteric buds, as demonstrated by Troma1 and GFP+ staining (Figures 4C–4E). Moreover, the immunostaining of day 3 organoids with nephron progenitor’s marker: Six2 presented integration of mESC-derived UB progenitors (GFP+) into the UB structure (Figure 4E). Together, this demonstrates that mESC-derived UB progenitors do integrate well into ureteric bud structures in chimeric renal organoids and are able not only to form UB *de novo* but also generate chimeric structures.

### Kidney organoids as models to study kidney development and drug toxicity

After transfer of the 3D chimeric organoids to organ culture, they self-organized to mature kidney organoid with proper nephron and collecting duct structures. As patterning of the nephron into its different segments begins at the renal vesicle stage during development (Georgas et al., 2009), we postulated that developmental patterning could be changed by chemical modulation of these endogenous signals.

Previous report revealed that Notch signaling is required for proximal tubule fate acquisition in the mammalian nephron (Cheng et al., 2007). We therefore treated the organoids with the Notch signaling inhibitor DAPT, as previous study shows that use of DAPT leads to suppression of proximal tubules formation in the human nephron organoid culture (Morizane et al., 2015, Cheng et al., 2003, Cheng et al., 2007). Thus, we added DAPT into the culture medium from day 3 to 11 of 3D organoid culture. Immunofluorescence analysis demonstrated that with the DAPT treatment, the formation of glomerulus and ureteric bud ducts is normal, but the proximal tubules were severely suppressed (Figure 5A).

**Figure 5.**
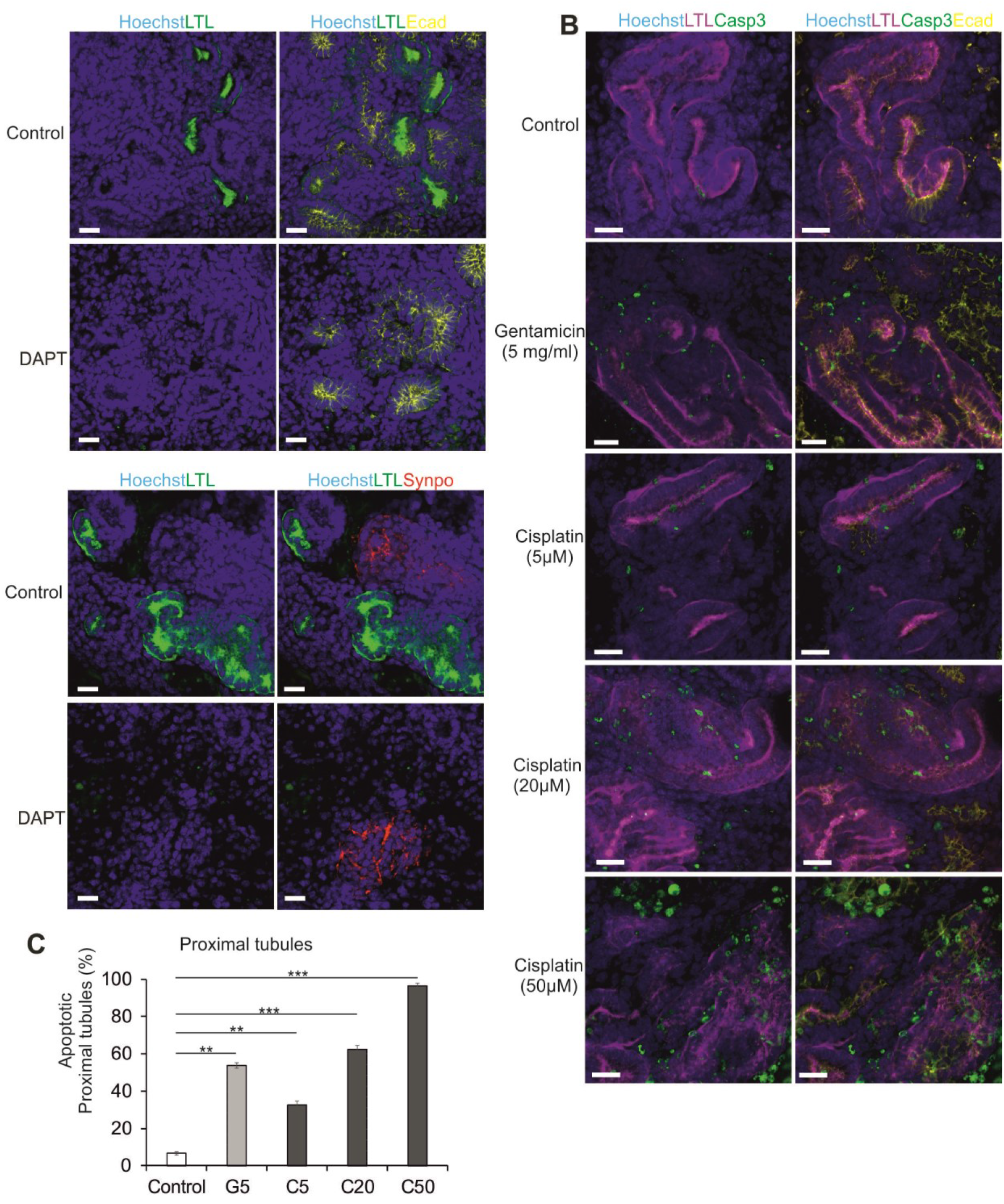
Kidney organoids models kidney development and injury. (A) Representative images of immunohistochemistry in kidney organoid presenting lack of UB tubular structures when treated with 10 mM DAPT from day 3 to 11. Notch inhibition suppressed the proximal tubule (LTL+) formation, but the UB (Ecad+) and glomerular (Synpo+) structures developed normally. n = 5; Scale bars, 20 μm. (B) Representative immunohistochemistry images of renal structures treated with gentamicin (5 mg/ml) from day 9 to 11 (48 hours) or cisplatin (5, 20, 50 μM) from day 10 to 11 (24 hours). Apoptotic cells were detected by cleaved caspase 3 antibody-staining (Casp3). n = 6; Scale bars, 20 μm. Quantification of the number of apoptotic proximal tubular cells in kidney organoids treated with gentamicin (G) (5 mg/ml) and cisplatin (C) (5, 20, 50 μM) at indicated doses. Data are expressed as mean ± s.e.m. (n=5); ** - p<0.01; *** - p<0.001.

In order to use stem cell derived kidney organoids for disease modelling and drug screening they need to present functional maturation of the nephrons within these organoids. To test whether these organoids could be used to study kidney injury and toxicity *in vitro*, we focused on drug nephrotoxicity, which have been shown as an important cause of acute kidney injury in hospitalized patients (Uchino et al., 2005). We treated the chimeric kidney organoids with gentamicin, a commonly used antibiotic with well-established proximal tubular toxicity, after 9 days of organ culture for 48 hours (Whiting and Brown, 1996). We also treated organoids with another nephrotoxicant, the cisplatin from day 10 for 24 hours. Cisplatin induces caspase mediated acute apoptosis of proximal tubular cells in the kidney (Mese et al., 2000). The whole-mount immunostaining with caspase3 of the control organoids showed random apoptotic interstitial cells; however, both gentamicin and cisplatin induced acute apoptosis in LTL+ proximal tubules (Figure 5B). The percentage of apoptotic proximal tubular cells induced by cisplatin increased in a dose-dependent manner; 50 μM cisplatin would lead to apoptosis of almost all of the proximal tubule cells (≈ 96%) (Figure 5C). We then immunostained the organoids with γH2AX, a DNA damage marker to distinguish between a generalized toxic effect and nephron segment-specific injury. Cisplatin at a dose of 5 μM upregulated γH2AX expression in LTL+ tubules only, whereas a higher dose of cisplatin (20 μM) resulted in more widespread γH2AX expression, consistent with more generalized cell toxicity (Figure S5).

To summarize, our work presents that the ESC-derived UB progenitors not only induced nephrogenesis in pMM, but also generated in that way chimeric renal organoids do present expected response to toxic chemicals and drugs. The established settings enable to exploit kidney organoid system to test the nephrotoxicity of drugs and other chemicals *in vitro*.

## DISSCUSSION

Kidney development starts from the interaction between the two precursors of the kidney, UB and MM. One major part of the MM cell population comprises the nephron progenitor cells (NPCs) which will differentiate into the nephrons; the ureteric bud will form the collecting duct tree. Protocols to direct differentiation of mouse and human pluripotent stem cells to renal progenitor cells and further self-organized kidney organoids containing nephrons have been well established (Takasato et al., 2015, Morizane et al., 2015, Freedman et al., 2015, Tan et al., 2018), but methods of differentiation of pluripotent stem cells specifically to ureteric bud progenitor cells are still not well developed (Xia et al., 2013, Taguchi and Nishinakamura, 2017).

Previous studies on generation ESC-derived UB, have shown the derivation of a UB-like population by selective induction with metanephric mesenchyme cells (Xia et al., 2013) or through CHIR99021 treatment for nephron differentiation of both ureteric bud and nephron structures (Takasato et al., 2015). However, the ESC-derived UB-like cells did not show the nephron progenitor induction (Xia et al., 2013), and therefore the inter-nephron connection by the collecting ducts was lacking (Takasato et al., 2015). Other group recently showed protocol to derive the UB structures from PSCs, and generated kidney organoids composed of mESCs-derived UB aggregated with primary MM, or mESCs-derived UB combined with mESC-derived NPC and primary stromal progenitor cells (SPs). Although successful, their protocol require to knock in markers into the PSCs, which leads to sorting of the specific marker expression cells at different differentiation stages. All this steps are therefore making this protocol technically complex (Taguchi and Nishinakamura, 2017). Here, we report the establishment of simple (directed with growth factors), efficient (>94% of Pax2+ cells) and reproducible differentiation protocol of mESC towards ureteric bud progenitor cells. These mESC-derived UB progenitor cells induced pMM cells to undergo nephrogenesis that led to development of well-structured nephrons. These nephrons consisted of glomeruli, proximal tubules, loops of Henle and distal tubules; and were connected with collecting duct generated by mESC-derived UB cells. Moreover, these mESC-derived UB progenitor cells formed the UB *de novo* when combined with pMM cells and generated chimeric structures when combined with the kidney rudiment cells (pUB and pMM) (summarized in Tables S1).

Previous studies showed that the ROCK kinases are VEGF downstream effectors, which negatively regulate the process of angiogenesis (Jens et al., 2009), we therefore added ROCK inhibitor to culture expecting increase in angiogenesis. However, we failed to see a difference between control and Y27 treated samples. Better results could be obtained by supplementing the medium with VEGF, which was already published by Freedman and co-workers, although they still did not observe endothelial cells entering the glomerular tuft (Czerniecki et al., 2018). Similarly, development of avascular glomeruli in organotypic kidney cultures and renal organoids was earlier reported by our research team (Halt et al., 2016). It seems that in absence of the blood flow, the endothelial cells are not able to properly interact with developing nephrons (Serluca et al., 2002) and formation of glomerular vasculature does not proceed further than migration of endothelial cells into vascular cleft region of an S-shaped stage nephron. It remains to be seen, if treatment with growth factors alone will enable vascularization of the glomeruli, or the renal organoids would have to be combined with blood vessels organoids (Reiner et al., 2019) to aid that process or providing the flow will be sufficient (Homan et al., 2019).

Even though organoids generated by mESC-derived UB progenitor cells did not contained patent vascular network, the developed nephrons did respond to toxicological tests as was expected. We and others (Morizane et al., 2015) show that the use of gentamycin and cisplatin induced apoptosis in proximal tubular cells. Thus, these organoids present a perfect platform to test nephrotoxicity of drugs.

In summary, we have developed an easy and reproductive protocol to generate UB progenitors from mESC. This work laid strong foundations to *in vitro* studies including disease modelling and drug discovery approaches; studies otherwise impossible to perform, or limited to the use of animal models and/or primary material that might not faithfully recapitulate developmental or disease progression features. Given the rapid progress in the field, we hope that in the near future, researchers would be able to generate full structure of nephrons in kidney organoids where the UB and MM parts would be derived from PSC. Using these cells should enable to generate not only well-structured nephrons, but also patent collecting duct tree. This would be a great first step for generating high throughput gene discovery models and advancing tissue engineering in producing organs for transplantations. However, these organoids would have to be successfully vascularized and grown to appropriate size. Nevertheless, these studies presented here, produce new insights into renal pathophysiology as well as open new avenues for developing new treatment options.

## EXPERIMENTAL PROCEDURES

The animal care and experimental procedures in this study were in accordance with Finnish national legislation on the use of laboratory animals, the European Convention for the protection of vertebrate animal used for experimental and other scientific purposes (ETS 123), and the EU Directive 86/609/EEC. The animal experimentation was also authorized by the Finnish National Animal Experiment Board (ELLA) as being compliant with the EU guidelines for animal research and welfare.

### Mouse ES cell line generation and maintenance

The mESC and mESC-GFP (pcDNA3.1 transfected) line was described previously (Tan et al., 2018) and maintained on mitotically inactivated mouse embryonic fibroblasts (MEFs) in Dulbecco’s modified Eagle’s medium (DMEM; Life Technologies) supplemented with 10% FBS (Gibco), 1% (vol/vol) nonessential amino acids (Life Technologies), 100 mM 2-mercaptoethanol (Nacalai Tesque), and 1,000 U/ml leukemia inhibitory factor (Millipore).

### Directed differentiation of mESCs to UB progenitors

Mouse ESCs were cultured as described above in MEF-coated 6-well plates in mESCs medium until reaching 70%-90% confluency. Mouse ESCs were passaged on 1% geltrex-coated 24 well plates at 30,000 cells/cm^2^ in 50% ReproFF2 (ReproCELL) – 50% CM (conditioned MEF medium) supplied with 10ng/ml FGF2 and 10ng/ml Activin A. After 2 days, cells reached 60–80% of confluency, we switched the medium to differentiation medium: the Advanced RPMI (Thermo Fisher Scientific) supplemented with 8 μM CHIR99021 and 4-8 ng/ml Noggin (R&D systems) for 2 days. Followed by treatment of the cells with Activin A (10 ng/ml) for 2 days and then 3 days with FGF9 (40 ng/ml) in Advanced RPMI medium.

### Gene expression analysis

An RNeasy kit (Qiagen) was used according to the manufacturer’s recommendations to extract the total RNA. cDNA synthesis (First Strand cDNA Synthesis Kit, ThermoFisher) was performed using standard protocols. qRT-PCR analyses were displayed with SYBR Green (Agilent) by an CFX96 Real-Time PCR machine. The Brilliant III SYBR^®^ Green QPCR Master Mix (Agilent Technologies) was used according to the manufacturer’s instructions. The *β-actin* probe served as a control to normalize the data. The gene expression experiments were performed in triplicates on three independent experiments. All the primer sequences are given in table S2.

### Flow cytometry

Differentiated cells were fixed with use BD Cytofix/Cytoperm™ solution (Cat. No. 554722) at 4°C for 20 min and then permeabilized by washing 2 x in 1 x BD Perm/Wash buffer (Cat. No. 554723). Cells were then resuspended in BD Perm/Wash buffer containing Pax2 primary antibody, incubated at 4°C for 30 minutes. We followed with washing cells 2x with 1x BD Perm/Wash buffer and resuspended in wash buffer containing mouse anti rabbit 647 second antibody. The we washed cells again 2 x prior to flow cytometry analysis.

### Chimera 3D kidney organoids formation

At day 9 of mESCs differentiation, which represents the UB progenitor cells stage, cells were dissociated into single cell suspension using TrypLE select (Life Technologies) after 3 x PBS washes, cells were reconstituted in organ culture medium. For the kidney organoids, we mixed dissected and dissociated pMM cells (from CD-1 pregnant females, at E11.5) as described before (Junttila et al., 2015) with differentiated UB progenitors at ratio 3:1, respectively. The cells were centrifuged for 4 min at 1380*g* to form a pellet (5 × 10^4^) in a Lo-Binding Eppendorf tubes. Following centrifugation, we carefully transferred the differentiated UB and pMM pellets to filter into Trowel culture to aggregate as an organoid. Changed the organ culture medium every 3-4 days.

For generation of whole kidney organoids we dissected mouse kidney rudiments at E11.5 from CD-1 pregnant females. Kidney rudiments were dissociated into single cell suspension as described before (Xia et al., 2013). After dissociation, the embryonic kidney cells (7 × 10^4^) were mixed with undifferentiated mESC or differentiated mESCs-derived UB progenitors (1 × 10^4^) to make the pellet. We then continued as described above.

### Whole-mount immunohistochemistry

The kidney organoids were washed 2 x with PBS and fixed with 100% cold Methanol (−20°C) for 30 min or 4% paraformaldehyde in PBS (organoid with GFP or dye) for 30 min. After fixation, organoids were washed at least 3 x in PBS and blocked in 0.1% Triton-X100, 1% BSA and 10% goat serum/0.02M glycine-PBS for 1-3 hours at room temperature. Incubation of the organoids with primary antibodies was performed in a blocking buffer overnight at 4°C. We followed with a wash 6x with PBS and incubated with secondary antibodies conjugated with Hoechst (Thermo Fisher Scientific) and Alexa Fluor 405, 488, 568, 546, or 647 (1:1000, Life technologies) and fluorescein anti-LTL (1:350, #FL-1321, Vector Laboratories) overnight at 4°C. Primary antibodies included Wt1 (1:100, #05-753, Millipore), Pax2 (1:200, #PRB-276P, Covance), Troma1 (1:200, DSHB), Gata3 (1:20, #AF2605-SP), E-cad (1:300, #610181, BD Biosciences), Synaptopodin (SYNPO) (1:4, #ABIN112223, antibody on line.com), Umod (1:25, #LS-C150268, LSBio), CD31 (1:100, #550274 BD Biosciences), Laminin (1:200, #L9393, Sigma), Cleaved Caspase-3 (1:200, #9661s, Cell Signaling Technology), γH2AX (1:200, #05-636, Millipore). The organoids were mounted with the Shandon™ Immu-Mount™ (Thermo Scientific™). A Zeiss LSM780 microscope and Zeiss Axiolab were used for image capture and analysis.

### Nephrotoxicity assay

3D kidney organoids were cultured in organ culture medium supplemented with gentamicin 5 mg/ml (Sigma, #G1264) for 48 h or cisplatin 5, 20 or 50 mM (Sigma, #P4394) for 24 h after day 8 of organ culture. Organoids were then fixed with 100% cold Methanol for 30 min for whole-mount immunohistochemistry.

## AUTHOR CONTRIBUTIONS

Z.T, A.R-R, I.S and S.J.V. designed the study; Z.T. performed experiments, made the figures and wrote the original draft, A.R-R and I.S revised the paper; all authors approved the final version of the manuscript.

## ACKNOWLEDGMENTS

We thank Paula Haipus, Hannele Härkman, Johanna Kekolahti-Liias and Sanna Kauppinen for technical assistance; Biocenter Oulu Transgenic core facility for the mouse ESCs; Dr. Jingdong Shan, Dr. Florence Naillat, MSc. Abhishek Sharma, Dr. Fariba Jian Motamedi and Prof. Andreas Schedl for the discussion. This work was supported financially by H2020 Marie Skłodowska-Curie Actions Innovative Training Network “RENALTRACT” (ID 642937), Academy of Finland (ID 315030, Centre of Excellence, ID 251314), Sigrid Jusélius Foundation, Cancer Research Foundation and Finnish Cultural Foundation (A.R-R).

## Supplemental Information

**Figure S1.**
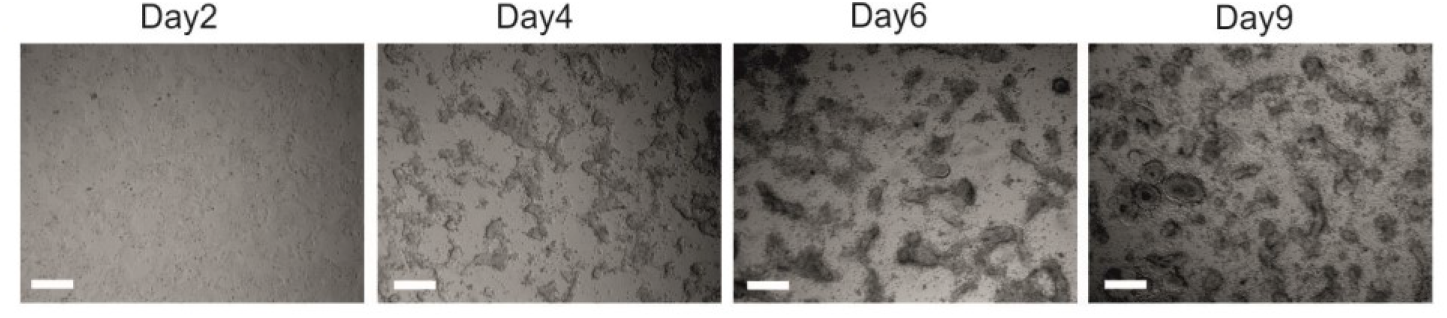
Bright field images of mESCs in monolayer (2D) cultures during differentiation into UB progentior cells. Consecutive days are shown. Scale bars, 200 μm.

**Figure S2.**
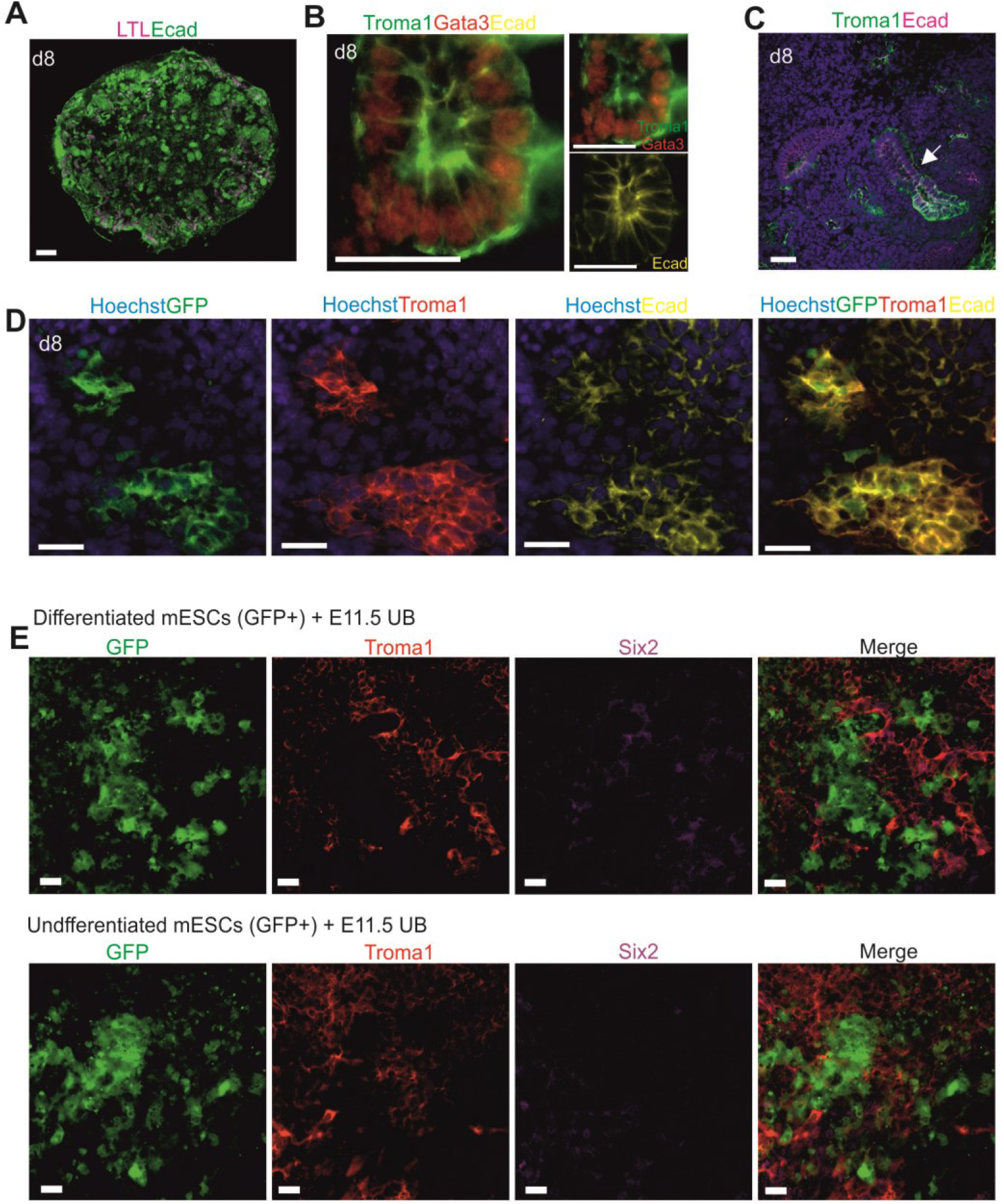
Differentiated mESCs induces embryonic MM to kidney organoids. (A) Tile scan immunofluorescence of a whole kidney organoid disp!aying tubular structural complexity. Scale bar, 200 μm. (B) Confocal imgae of collecting duct structure. Scale bare 20 μm. (C) Fused collecting ducts in the organoids. Scale bar, 20 μm. (D) Reconstruction kidney organoids with Troma1 +Ecad+ collecting ducts are derived from GFP+ mESCs. Scale bars, 20 μm. (E) Confocol images showed no visible signs of *in vitro* nephrogenesis when purified embryonic ureteric bud (UB) cultures with differentiated or undifferentiated GFP+ mESCs. Scale bars, 20 μm.

**Figure S3.**
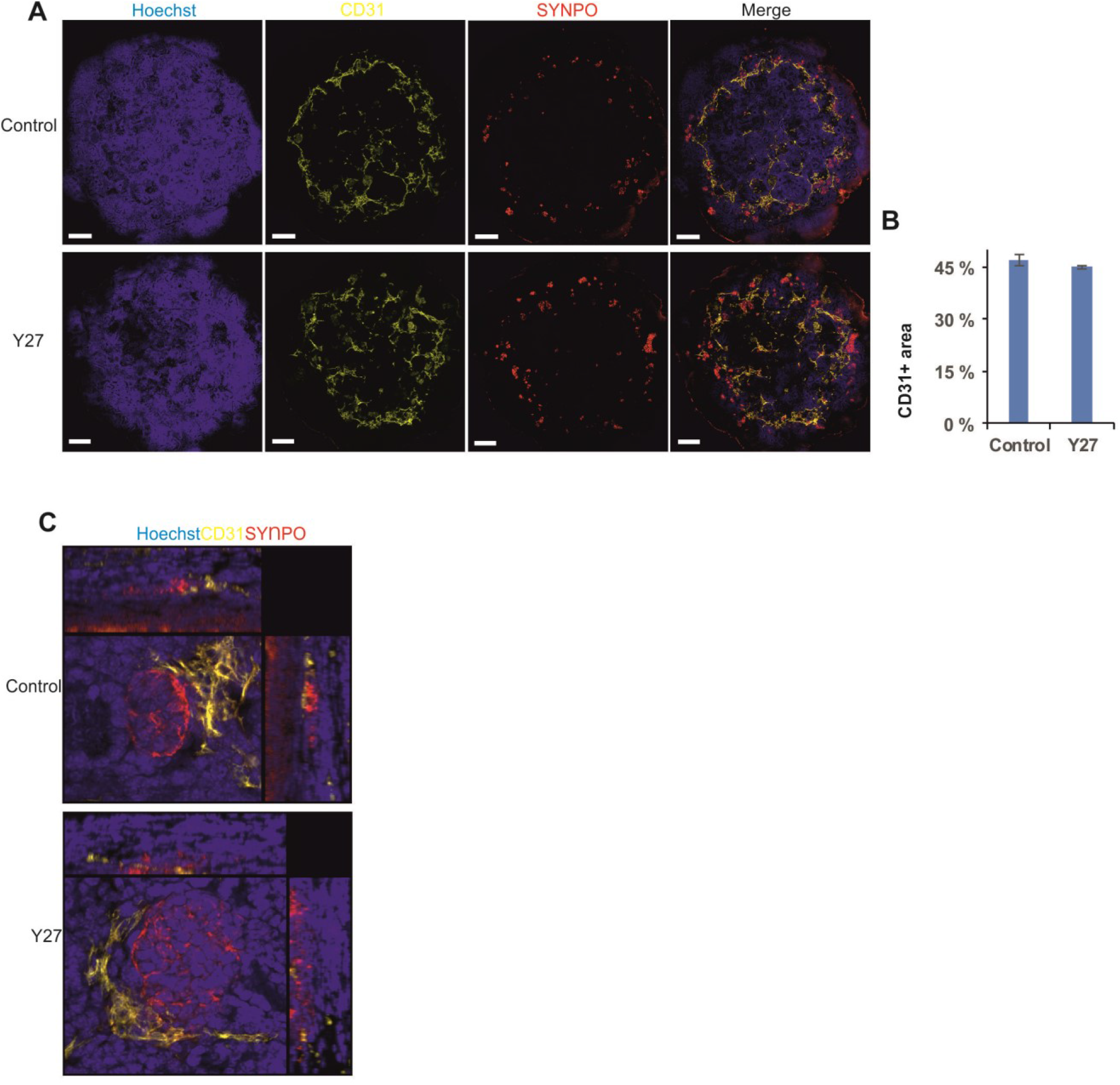
Vascularization of the kidney organoids. (A) Tile scan immunofluorescence of the whole kidney organoids displaying vascularization network treated with Rock inhibitor (Y27) or without (Control). Scare bars, 2Ũ0 μm. (B) Percentage of the total culture area occupied by cells expressing CD31, n=6 experiments (+/− S.E.M.)· (C) Confocal multiple image projection samples showing glomeruli (Synpo+) with endothelial cells (CD31+) in organoids treated with Y27 or without treatment (control). Scale bar, 20 μm.

**Figure S4.**
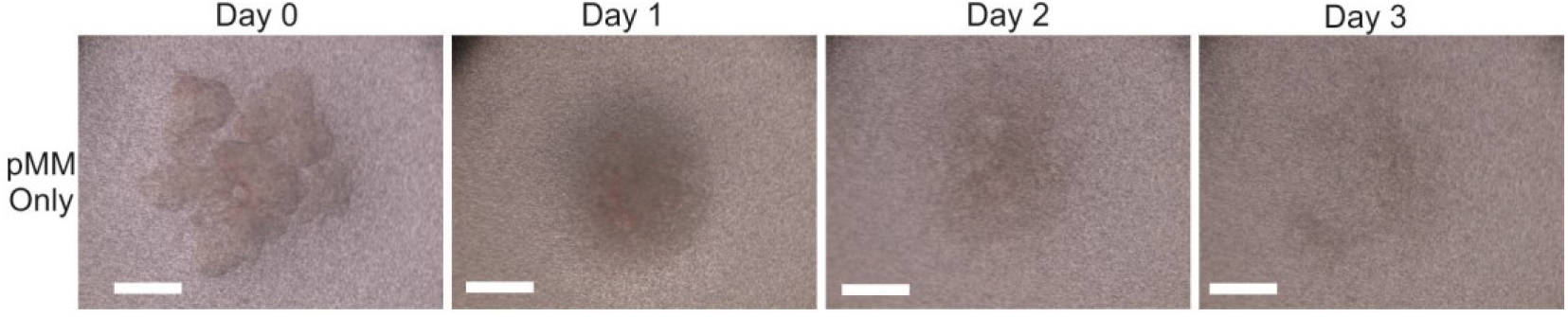
Only primary MM (pMM) on the 3D Trowell culture died after 3 days (n=3). Scare bars, 500 μm.

**Figure S5.**
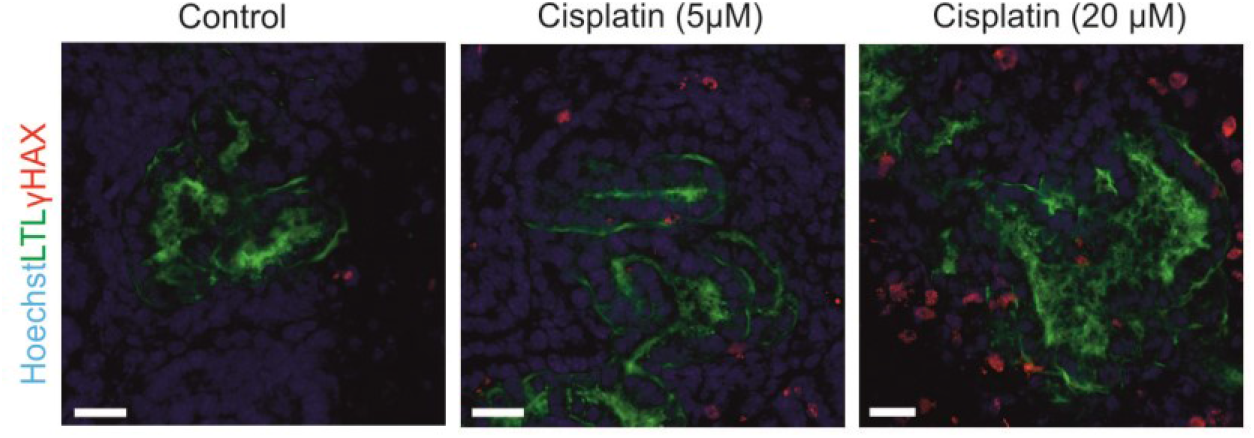
Representative immunohistochemistry of organoids treated with cisplatin (5 or 20 μM) from day 10 to 11 (24 h) present DNA damage, n = 5. Scale bars, 20 μm.

**Table S1.** Summary of available protocols of differentiation of mouse and human PSC to UB lineages.

|  |  | Our protocol | Xia et al., 2013 | Takasato et al., 2015 | Taguchi et al., 2017 | Taguchi et al., 2017 |
| --- | --- | --- | --- | --- | --- | --- |
| Cell type |  | mESC | hESC/hiPSC | hESC/hiPSC | mESC | hiPSC |
| Differentiation condition | Basic differentiation medium | Advance RPMI 1640 | DMEM/F12 | APEL | IMDM/F12 | IMDM/F12 |
|  | Specific factors (PSC differentiated to IM) | Activin A | BMP4 | CHIR99021 | Activin A | Activin A |
|  |  | FGF2 | FGF2 | FGF9 | BMP4 | BMP4 |
|  |  | CHIR99021 | Retinoic Acid |  | CHIR99021 | CHIR99021 |
|  |  | Noggin | Activin A |  | Retinoic Acid | Retinoic Acid |
|  |  |  | BMP2 |  | FGF9 | FGF9 |
|  |  |  |  |  | SB431542 | LDN193189 |
|  |  |  |  |  |  | SB431542 |
|  | Specific factors (IM differentiated to UB) | FGF9 |  | CHIR99021 | Retinoic Acid | Retinoic Acid |
|  |  |  |  | FGF9 | CHIR99021 | CHIR99021 |
|  |  |  |  |  | FGF9 | FGF9 |
|  |  |  |  |  | Y27632 | Y27632 |
|  |  |  |  |  | GDNF | GDNF |
|  |  |  |  |  |  | LDN193189 |
|  |  |  |  |  |  | FGF1 |
| FACS sorting step |  | no | no | no | yes, FACS sorting of the Cxcr4+/Kit+ cells | yes, FACS sorting of the CXCR4+/KIT+ cells |
| Generation of UB *de novo* |  | yes | no | yes | yes | yes |
| Collecting duct connect with nephron tubule |  | yes | not shown | yes | yes | not shown |
| Integration of UB cells |  | yes, Integrated into E11.5 mouse UB on organ culture | yes, Integrated into E11.5 mouse UB on organ culture | not shown | not shown | not shown |

**Table S2.**
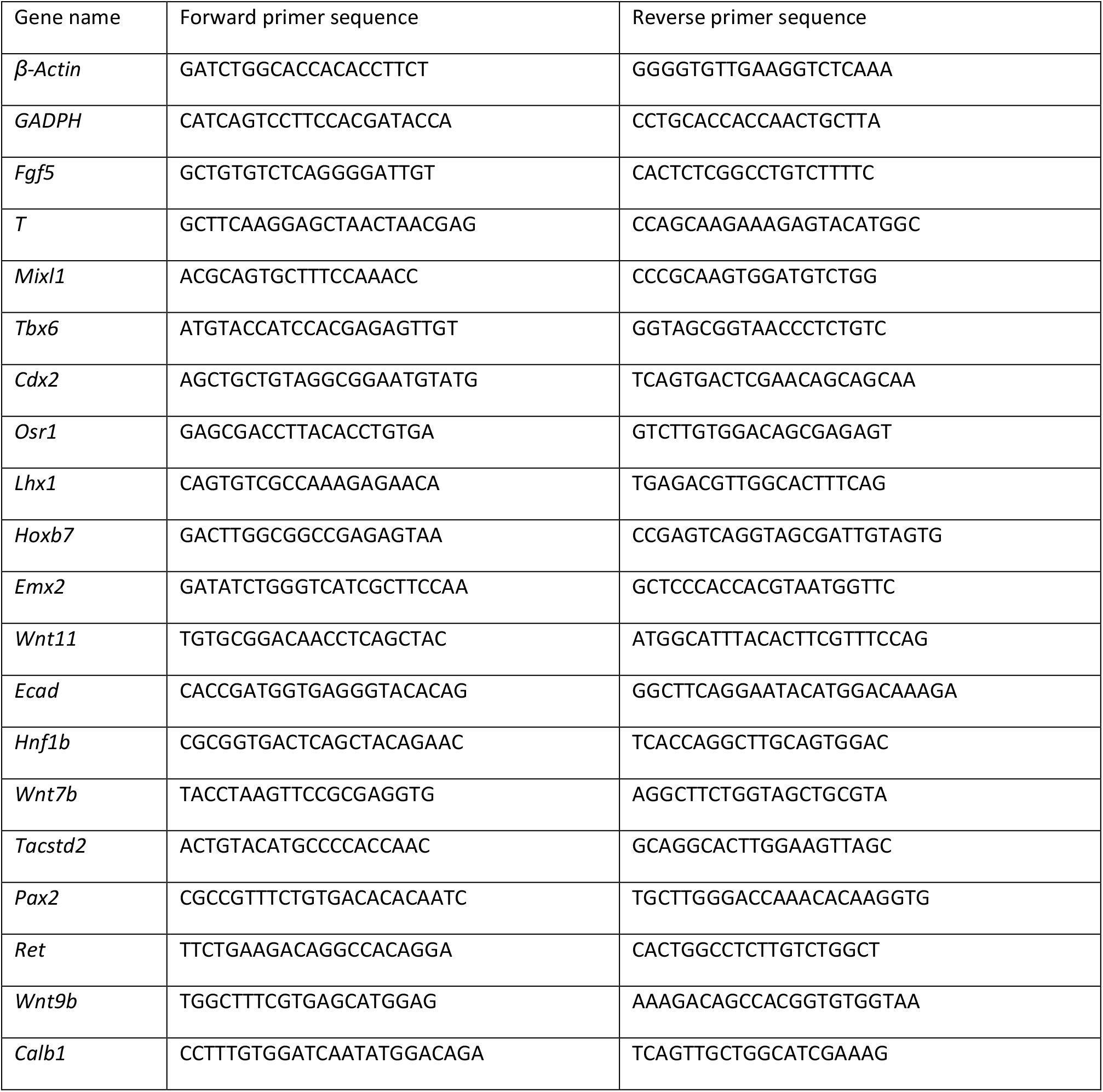
Primers (5’-3’) used in the study.

